# Efficient Method for Estimating the Frequency of Plasmidless Cell Generation per Cell Division

**DOI:** 10.64898/2026.11.16.766249

**Authors:** Christina Egami, Viraj Arora, Manel Camps

**Affiliations:** Department of Microbiology and Environmental Toxicology, University of California at Santa Cruz, Santa Cruz, CA 95064, USA

## Abstract

Plasmids are self-replicating pieces of DNA that play a major role in the evolution and spread of adaptive genes in bacteria. Plasmids tend to form stable associations with their hosts, including plasmids such as small, multicopy plasmids that typically lack specialized partition and horizontal transfer systems. Using a ColE1 plasmid model, we present a method that quantifies the frequency of plasmid loss over a short, defined time window. This method is based on the transient survival of plasmidless cells in the presence of selecting antibiotic and it has two main advantages over plasmid visualization approaches: (1) it does not require modifications in plasmid sequence, preventing artifactual results linked to sequence alteration, and (2) it can unambiguously resolve plasmidless cells relative to cells with a low plasmid copy number. Thus, our approach is ideal for the study of the stability of naturally-occurring small, multicopy plasmids or for the identification of modulators of stability. Using our method, we describe a mutant whose long-term stability is exclusively dependent on increased accurate segregation. Finally, we report that moderate sequence manipulations (outside *ori* sequence) do not have a major impact on early plasmid loss, which is relevant for the design of plasmid tagging approaches.

## Introduction

Plasmids are autonomously-replicating pieces of DNA that are ubiquitous in bacteria ^1^. In addition to the genes needed for reproduction, maintenance and transmission, most plasmids encode “accessory” genes. This plasmid-borne accessory cargo gene can provide benefits to the host by expanding their ecological niche, by degrading toxic compounds or providing new metabolic capabilities ^2-4^. Plasmids are also widely used as tools for the production of proteins, enzymes, antibodies, drugs and other organic compounds in large-scale fermentation processes ^5,6^.

At the same time, plasmids also cause fitness costs to their hosts, particularly when introduced in a new bacterium ^7-11^. These costs are associated with genes needed for plasmid reproduction, transmission and maintenance, which can be considered selfish (parasitic) because they don’t necessarily share evolutionary interests with the chromosome ^2,12^ or are due to genetic conflicts between the plasmid or its products and the host’s DNA, RNA or proteins ^13^. Thus, plasmid-free cells have a considerable fitness advantage over plasmid-bearing cells. Other factors affecting the long-term stability of plasmids include the rate of horizontal transfer by conjugation, founder effects acting on plasmid-bacterium associations, and the presence of specialized plasmid maintenance systems such as active partitioning or post-segregational killing systems ^14-16^. Therefore, in the absence of specialized maintenance systems, theory predicts the complete loss of a plasmid from a bacterial population by purifying selection unless a positive selection for plasmid traits is present or the plasmid is able to spread horizontally fast enough to ensure survival as a pure genetic parasite. In addition, consistent positive selection for plasmid-borne beneficial traits should lead to their integration into the chromosome, rendering the plasmid dispensable. In view of these considerations, the observed long-term associations between plasmids and their hosts are hard to explain, a phenomenon known as “the plasmid paradox ^11,17,18^.

The long-term maintenance of small, multicopy plasmids is particularly paradoxical, as these plasmids typically lack specialized maintenance or complete conjugative systems yet they are extremely prevalent in the environment ^19^ and in the clinic ^19,20^ The long-term stability of these plasmids is assumed to be dependent on the frequency of spontaneous generation of plasmid-free cells ^21^. The simplest model assumes a random distribution of plasmids in the mother cells and random segregation during cell division. According to this random distribution model, the probability of plasmidless daughter generation is P_0_=2 ^(1-n)^, where n is the plasmid copy number of the mother cell ^22^. Using advanced quantitative methods, Lau et al. documented random fluctuations in plasmid copy number in the population that increase the probability of random plasmid loss relative to estimates based on average plasmid copy number ^23^. In addition, the number of independently segregating plasmid units can be reduced by the formation of multimers ^24^ or by the formation of clusters or “foci” ^25,26^, increasing the probability of plasmid loss.

Thus, in the absence of specialized maintenance systems or horizontal transfer, the generation of plasmid-free cells can be understood as resulting from an equilibrium between plasmid replication, foci formation, multimer resolution and partition. Here we describe a method to measure the frequency of spontaneous plasmidless cell generation in liquid culture to facilitate the study of plasmid stability **(Fig. 1)**. As a model system we use a small multicopy plasmid with a ColE1-like origin of replication, an antibiotic selectable marker, and green fluorescent protein (GFP) as a reporter gene. Our method is based on the observation that plasmidless cells can remain transiently viable in liquid cultures in the presence of antibiotic selection reported in one of our earlier studies ^27^. This approach allows us to measure early plasmid loss, *i*.*e*. the plasmid loss that occurs during a defined window of time that is short enough to make the contribution of differential fitness negligible. We then convert the results to plasmid loss events per cell division using a mathematical model that integrates the time window and growth rate.

**Figure 1.**
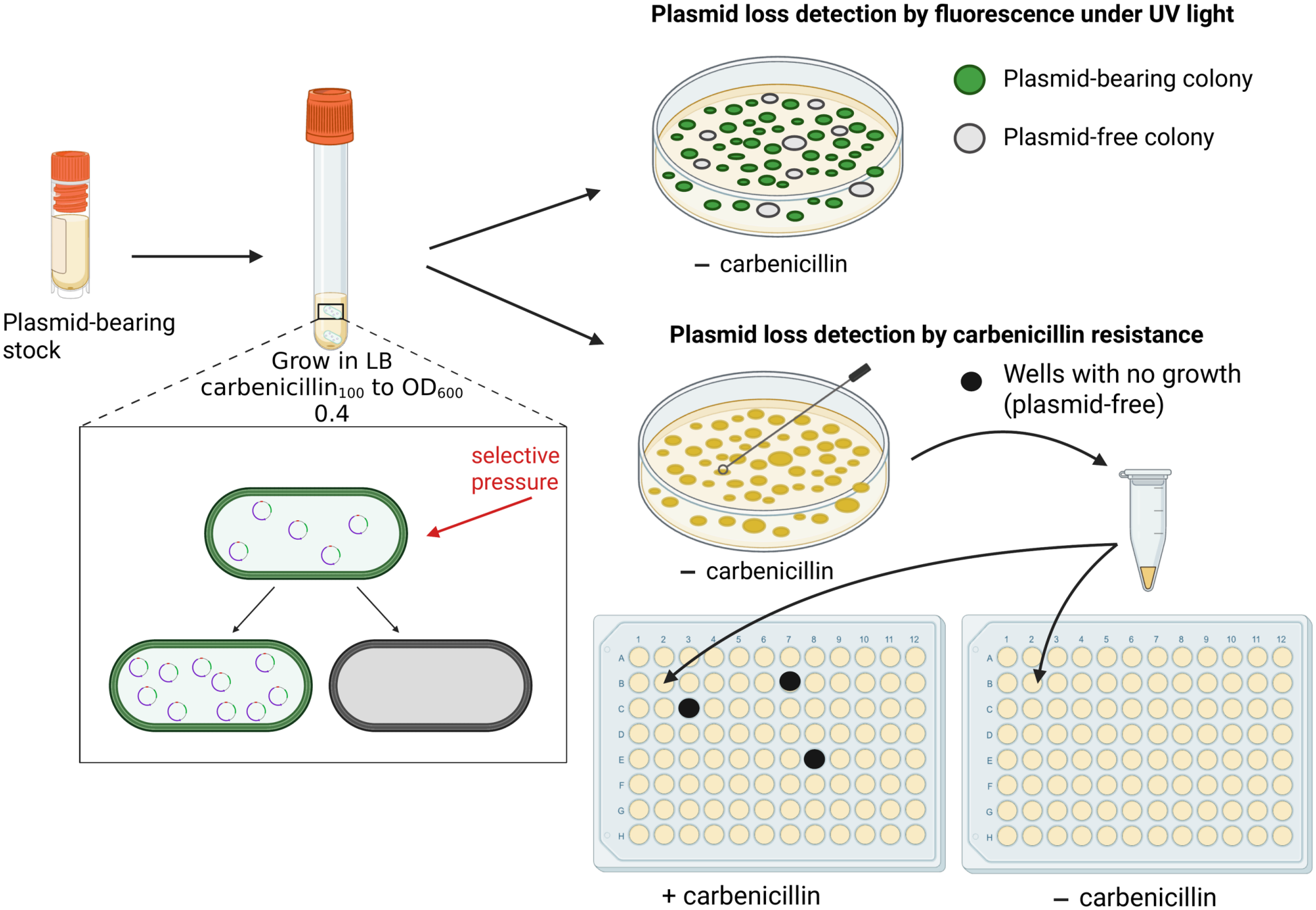
Method for early plasmid loss detection using loss of fluorescence or of carbenicillin resistance as readouts. Cells carrying the plasmid are grown in LB with carbenicillin to OD_600_ ≈ 0.4, maintaining selective pressure for the plasmid; during growth, inaccurate segregation gives rise to plasmid-free daughter cells. Loss is then scored on non-selective (carbenicillin-free) plates using one of two markers present in the plasmid. (Top) Loss of fluorescence: colonies are imaged under UV, where plasmid-bearing colonies exhibit fluorescence (and appear as bright green) and plasmid-free colonies do not (and remain dark). Cycle 3 GFP can in principle be replaced by another fluorimetric or colorimetric marker. (Bottom) Loss of carbenicillin resistance: individual colonies from a plain LB Petri dish (i.e grown without added carbenicillin) are picked into suspension and arrayed into paired +carbenicillin and −carbenicillin 96-well plates; plasmidless colonies can be detected as growing in liquid media but fail to grow in media supplemented with carbenicillin. Carbenicillin can be replaced by another selection so long as the plasmid carries the corresponding selectable marker.

The approach that we describe here has considerable advantages over visualization methods and it can advance our understanding of long-term multicopy plasmid stability, which is a question of considerable ecological and biotechnological relevance.

## Results

### Experimental System for the study of determinants of multicopy plasmid stability

Our experimental model consists of two versions of a multicopy plasmid, pGFPuv and pGFPck, which are described in detail in ^28^. They contain pUC19 (a ColE1-like plasmid origin of replication), TEM-1 β-lactamase as a selectable marker, and cycle 3 GFP as a fluorescent reporter; pGFPck additionally carries a kanamycin resistance gene. As a highly-stable plasmid control, we also included a superstable pGFPuv mutant, which has two mutations that synergistically confer our pGFPuv plasmid with a high level of long-term stability ^27^.

As a host we use Top10 *E. coli* cells, a strain used for recombinant gene expression. This strain has a *recA1* allele, which exhibits decreased RecA’s ATPase activity in the presence of single-stranded DNA, resulting in reduced rates of genetic recombination ^29,30^. Suppressing homologous recombination is important to minimize concatemer formation, which is known to contribute to reducing the number of effective partitioning units during cell division ^24,31^.

In our simplified experimental model system, we assumed the long-term stability of the plasmid to be dependent mainly on two factors, namely the rate of segregation of plasmidless cells by asymmetrical plasmid segregation, and the fitness cost of the plasmid on the host. We first measured the impact of pGFPuv WT and DAS G512A on the hosts’ fitness by looking at their growth rates **(Fig. 2a)**. Both pGFPuv WT and DAS G512A mutant exhibited similar decreases in growth rate, which is surprising given the much stronger stability exhibited by the DAS G512 plasmid, as established by an earlier study from our group [27]. We also measured the plasmid copy number of both plasmids by qPCR and found no significant differences between pGFPuv WT and DAS G512A, (66.7 and 75.6 copies per chromosome, respectively; **Fig. 2b**). Thus, the higher stability of DAS G512A cannot be attributed to differences in either fitness cost or plasmid copy number.

**Figure 2.**
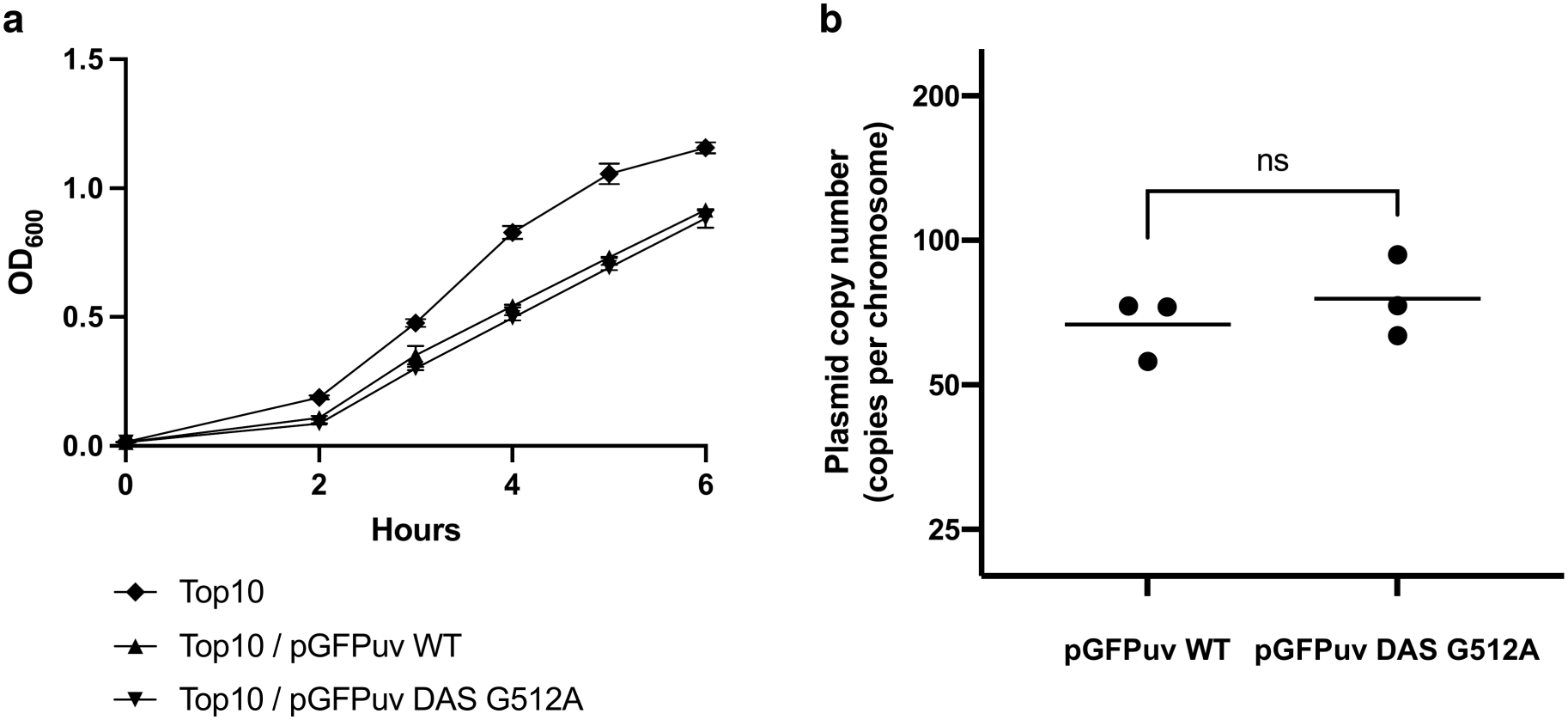
Fitness cost and copy number of pGFPuv WT and the DAS G512A variant. **(a)** The growth of plasmid-free Top10 and Top10 carrying the indicated plasmids was monitored by OD_600_ over 6 h. Top10 was grown in LB; plasmid-bearing strains were grown in LB supplemented with 100 μg/ml carbenicillin. Points are mean ± s.d., n = 3 biological replicates. The slower growth of plasmid-bearing strains reflects the fitness cost of plasmid carriage. **(b)** Plasmid copy number (copies per chromosome) of pGFPuv WT and DAS G512A, determined by qPCR. Points are individual biological replicates (n = 3) and horizontal lines indicate the geometric mean. pGFPuv WT and DAS G512A did not differ significantly (ns, adjusted P = 0.77). Copy numbers of all four constructs (pGFPuv WT, DAS G512A, pGFPck WT and Δ6; see Fig. 8b) were measured in the same experiment and analyzed together by Welch’s one-way ANOVA on log-transformed values (P = 0.0014) with Dunnett’s T3 multiple comparisons test for three planned comparisons.

### Detection of viable plasmidless cells in culture in the presence of antibiotic

As a proxy for detecting individual asymmetrical plasmid segregation events producing plasmidless cells, we sought to quantify plasmid loss within a window of time close to a single replication cycle. In a previous article, we noted that plasmidless cells generated from cells bearing our pGFPuv plasmid can transiently survive in the presence of carbenicillin selection in sufficient numbers to be easily detectable by plating ^27^. In that report, plasmid loss was detected in three ways, namely by loss of GFP fluorescence in single cells run through FACS, by loss of colony fluorescence, and by loss of carbenicillin resistance ^27^.

To confirm those results, we plated a countable number of colony-forming units (CFUs) from WT pGFPuv-bearing cultures grown in exponential phase in the presence of carbenicillin on LB plates. The GFP^lo^ colonies detected on these plates represented 1.45% of the total (**Fig. 3a, b, c)**. We confirmed that GFP^lo^ colonies indeed grew out from plasmidless single cells by growing 60 randomly selected GFP^lo^ colonies in the presence and absence of carbenicillin. All 60 picked colonies grew in LB without carbenicillin, showing that we successfully transferred viable cells from the colony to our media, but none grew with added carbenicillin, confirming that these colonies harbored no plasmid. As a control, we also grew 60 GFP^pos^ colonies and they all grew both in the presence and absence of carbenicillin, confirming that GFP^lo^ failure to grow in carbenicillin was not due to inconsistent cell transfer. By contrast, in cultures bearing the DAS G512A pGFPuv plasmid, we saw no GFP^lo^ colonies (out of a total of 2044), indicating an early plasmid loss frequency in culture of less than <0.05% (with a 95 percent confidence) (**Fig. 3a**). These experiments confirmed that in our experimental model system, colony fluorescence aligns perfectly with carbenicillin resistance and that either reporter can be used to track the presence or absence of plasmids in individual cells.

**Figure 3.**
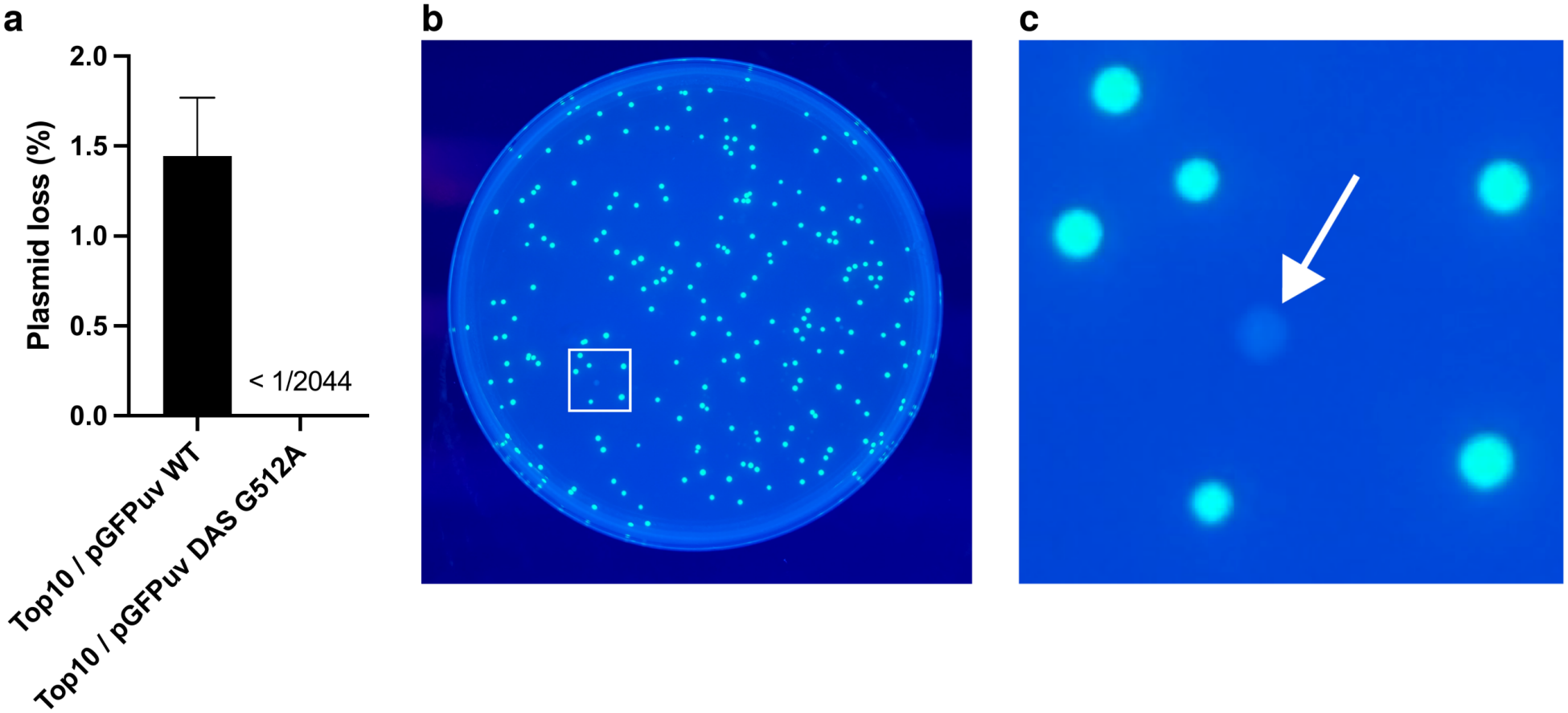
Plasmid loss detection. Early plasmid loss was measured for Top10/pGFPuv WT and the DAS G512A variant. **(a)** Loss frequency scored by loss of colony fluorescence, plotted as % loss (mean ± s.d., n = 3 biological replicates): WT = 1.45% (26 GFP^lo^ /1789 colonies), DAS G512A below detection (<1/2044). **(b)** Representative LB plate imaged under UV, from which counts were obtained. **(c)** Digital enlargement of the boxed region in (b) — a cropped detail of the same image. Arrow marks GFP^lo^ (plasmid-free) colonies.

### Establishing the window of transient survival in the presence of antibiotic selection

We reasoned that the survival of plasmidless cells in the presence of antibiotics must correspond to recent plasmid losses, *i*.*e*. to cells that had not yet been killed by the antibiotic once they lost their plasmid. Indeed, the antibiotic used as a selection, carbenicillin, is known to kill by a delayed process involving a weakening of the cell wall culminating in cell lysis ^32,33^. If our interpretation is true, this provides a defined time window for the plasmid loss, allowing us to estimate the fraction of cells that lose their plasmid at each cell division cycle by factoring in growth rate.

We established the window during which plasmid-free cells can still survive in the presence of antibiotic exposure by exposing a plasmidless cell culture to carbenicillin. In parallel, we include plasmid-bearing cells as antibiotic resistant controls. We did this in liquid culture to mimic our experimental system as closely as possible and to prevent carbenicillin-sensitive cell survival through the release of free beta-lactamase into the media by carbenicillin-resistant cells, which is known to happen in solid media ^34^. To shorten the time-window of survival further, we also treated cultures with kanamycin, which induces cell death faster through inhibition of protein synthesis and the generation of reactive oxygen species ^35^.

Changes in cell density in culture due to continued growth and/or to cell lysis were monitored by OD_600_. **Fig. 4a** shows the time-course following drug exposure to carbenicillin and kanamycin measured by OD_600_. Carbenicillin did not appear to have an immediate effect on growth, allowing an increase in OD_600_ for another ∼30 minutes after initial exposure, arresting and even losing cell density after this time point. This is consistent with carbenicillin’s mechanism of action (targeting the cell wall, a cell structure that stabilizes cells against osmotic stress) and its classification as a β-lactam antibiotic belonging to group 1, which has no effect on doubling time and leads to formation of filamentous cells through inhibition of septation, followed by lysis ^33^. Kanamycin, by contrast, arrests growth earlier (at ∼20 min) and does not lead to a substantial loss in cell density. **Fig. 4b** shows that in the presence of the pGFPck plasmid, the addition of carbenicillin or kanamycin does not have a significant effect on growth, confirming that the effects of drug treatment seen in the absence of the plasmid in panel **a** can be solely attributed to the loss of the plasmid.

**Figure 4.**
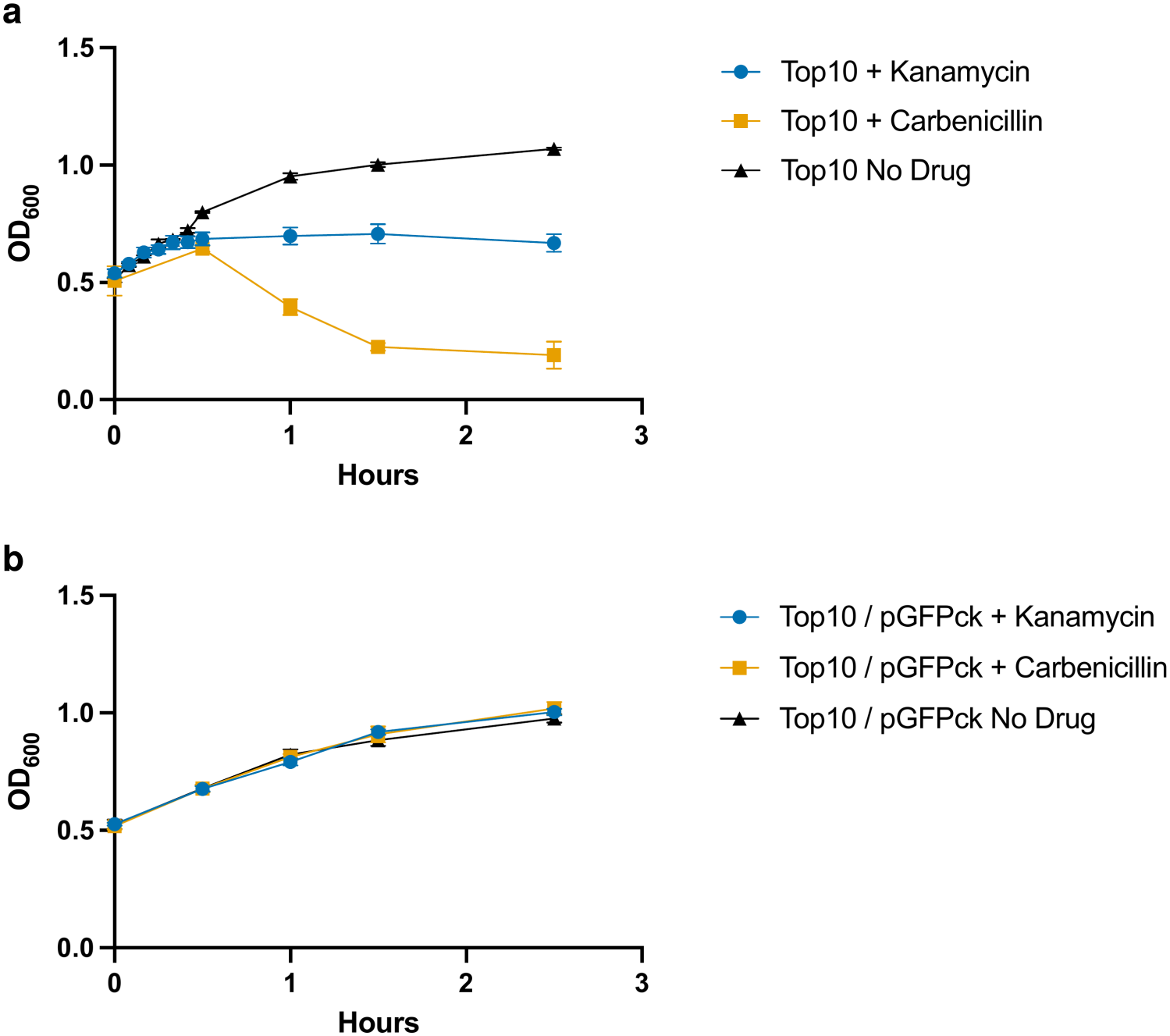
Effect of carbenicillin on the growth of plasmid-free and plasmid-retaining cells (OD600). Top10 cell cultures, either plasmid-free or carrying pGFPck, were exposed to carbenicillin, kanamycin, or no drug and followed by OD_600_ over 2.5 h. Points are mean ± s.d., n = 3 biological replicates. **(a)** Plasmid-free Top10: the no-drug culture continues to grow, kanamycin plateaus at ∼20 min, and carbenicillin declines after ∼0.5 h. **(b)** Top10/pGFPck: all three cultures grow indistinguishably, indicating that neither drug arrests the growth of plasmid-retaining cells.

Loss of viability was measured by vital staining and plating efficiency (PE) (**Fig. 5-6**). The vital staining consists of two nucleic acid stains, propidium iodide (PI) and SYTO 9. SYTO 9 is a membrane-permeable, green fluorescent dye. PI emits red fluorescence and is membrane impermeable and is therefore excluded by “healthy” cells. In dead or dying cells, since PI has a stronger affinity for nucleic acids than SYTO 9, SYTO 9 will be displaced from nucleic acids and the cell(s) will fluoresce in red. The results are shown in **Fig. 5a**. Carbenicillin exhibited delayed cytotoxicity, with the proportion of live-staining cells remaining essentially unchanged during the first 30 min of exposure (87.4% at 30 min, compared to 88.2% in the untreated culture) before decreasing to approximately 22.4% at 1 h. Representative fields of cells treated with carbenicillin stained with PI and SYTO 9 (merged) are shown in **Fig. 5c, d** (30 min, 1h). After 1h of carbenicillin exposure, microscopy shows filament formation (white arrows) and a substantial reduction in cell density (i.e. # cells per field) (**Fig. 5b**). This observation is consistent with the drop in OD values below values seen prior to drug exposure and is indicative of extensive carbenicillin-induced lysis ^33^. In contrast, kanamycin rapidly reduced the proportion of live-staining cells, decreasing from approximately 88% at 0 min to 48% within 15 min and to only 4% by 30 min. Cells exposed to kanamycin for 30 min and 1h are shown in **Fig. 5e, f**, respectively. In this case, we can see a much larger proportion of dead (red) cells and no substantial change in cell density over time.

**Figure 5.**
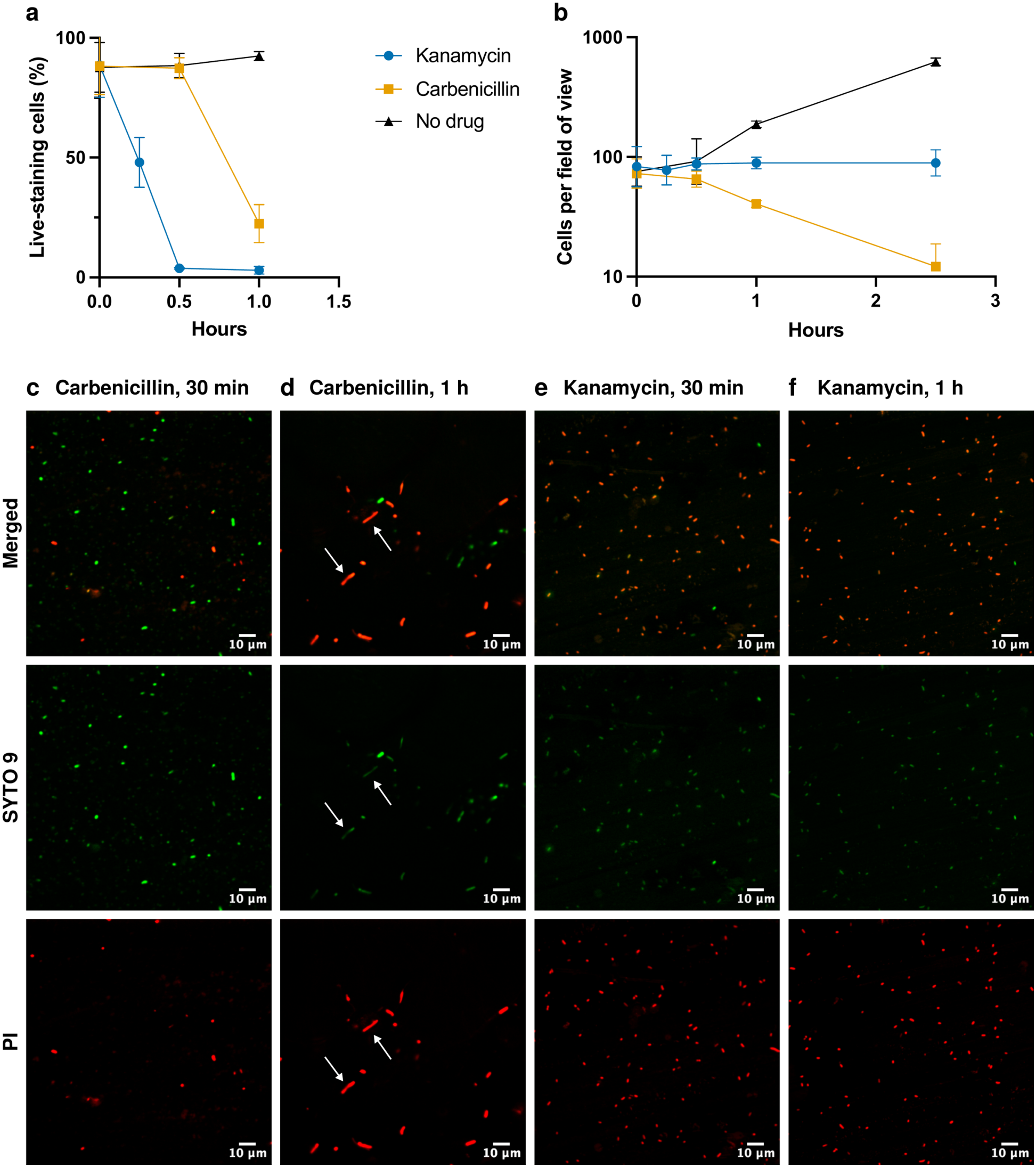
Effect of carbenicillin and kanamycin on plasmid-free cell viability, as seen by vital staining. Plasmid-free Top10 cells were exposed to no drug (black triangle), to carbenicillin (orange square), or to kanamycin (blue circle). **(a)** Quantification of live-staining cell fraction (%) over 1 h of exposure using merged SYTO 9/PI staining. Points are mean ± s.d., n = 3 biological replicates. **(b)** Cell density quantification over 2.5 h, quantified as cells per field of view; points are mean ± s.d. of three independent samples per condition. Because carbenicillin induces filamentation (panel d), the decline under carbenicillin reflects both cell lysis and the scoring of individual filaments — each comprising several cells — as single objects. **(c-f)** Representative fields imaged with a 63× oil-immersion objective; each column shows the merged image, SYTO 9 (green) and PI (red) channels of the same field. **(c)** 30 min carbenicillin. **(d)** 1 h carbenicillin; filaments marked by arrows. **(e)** 30 min kanamycin. **(f)** 1 h kanamycin. Scale bars, 10 µm.

Finally, **Fig. 6** shows survival to drug exposure as seen by PE. The results follow a similar trend as those seen with vital staining, with kanamycin killing much faster than carbenicillin. However, PE is a considerably more sensitive readout. After 30 min of carbenicillin exposure, we already see a 48% loss in colony forming units, compared to only ∼1% loss of vital staining at this same time point. Similarly, 15 minutes of kanamycin exposure already resulted in 98.5% loss in colony forming units, compared to only 52% staining red (dead) at the same time point.

**Figure 6.**
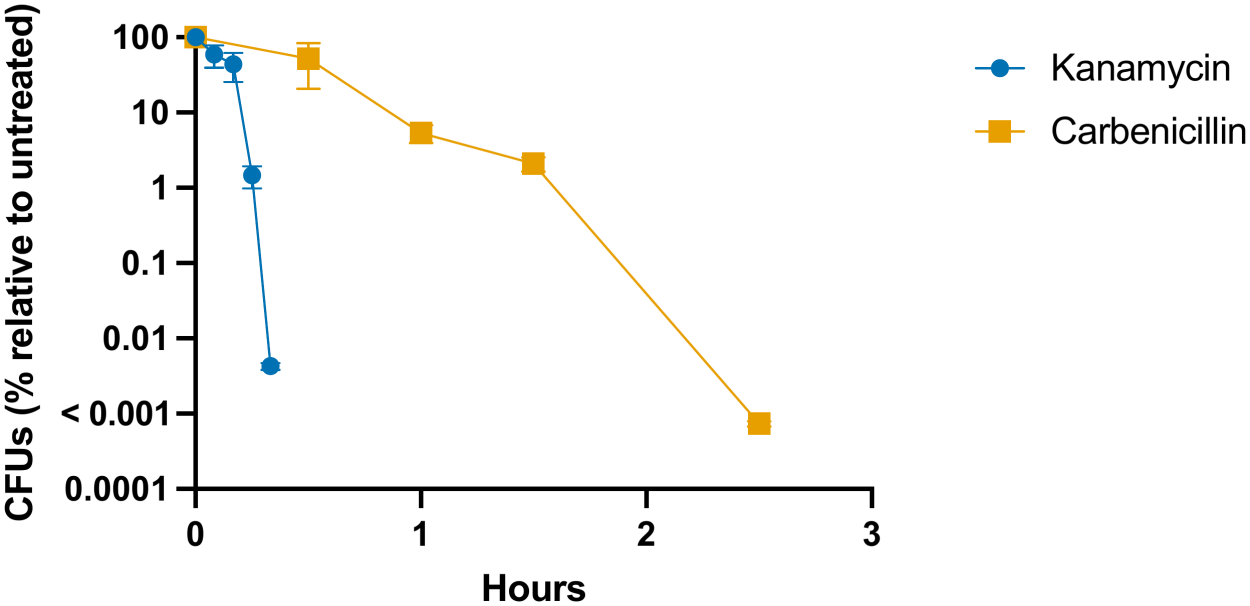
Time-course of plating efficiency of plasmid-free cells exposed to carbenicillin or kanamycin. Plasmid-free Top10 cells were exposed to carbenicillin (orange squares) or kanamycin (blue circles), and viable counts were determined at intervals and expressed as CFU as a percentage of the untreated control (log scale). Points are mean ± s.d., n = 3 biological replicates.

### Early plasmid loss detection using kanamycin

Carbenicillin and kanamycin exhibit considerable differences in the kinetics and mechanism of cytotoxicity. However, we reasoned that if our method is indeed measuring the frequency of spontaneous plasmid loss, the estimates we obtain should be consistent regardless of the drug used. To determine the percentage of plasmidless cells found in liquid cultures selected with kanamycin, we had to use a pGFPuv plasmid derivative, pGFPck, which has the *nptII* gene conferring kanamycin resistance

To our surprise, we saw that pGFPck plasmid exhibited a strong enhancement in long-term stability (**Fig. 7a**). This dramatically increased the number of colonies needing to be screened to detect early plasmid loss. To address this limitation, we attempted to destabilize the pGFPck plasmid by generating eight microdeletions spanning the length of the plasmid sequence (**Fig. 7b**). The results are shown in **Fig. 7c**. Six of the eight microdeletions showed no detectable early plasmid loss in the presence of carbenicillin screening comparable numbers of colonies. A microdeletion between the kanamycin resistance gene and the β-lactamase gene (Δ3), however, showed one plasmidless colony out of 1173 screened colonies, suggesting a mild impact on short-term plasmid stability but a larger sample size would be needed to confirm this conclusion. The microdeletion that precedes the ColE1 *ori* (Δ6) shows a clearer destabilization effect, with 8 out of 1027 plasmidless colonies (at least a ∼8.5-fold increase relative to WT pGFPck).

**Figure 7.**
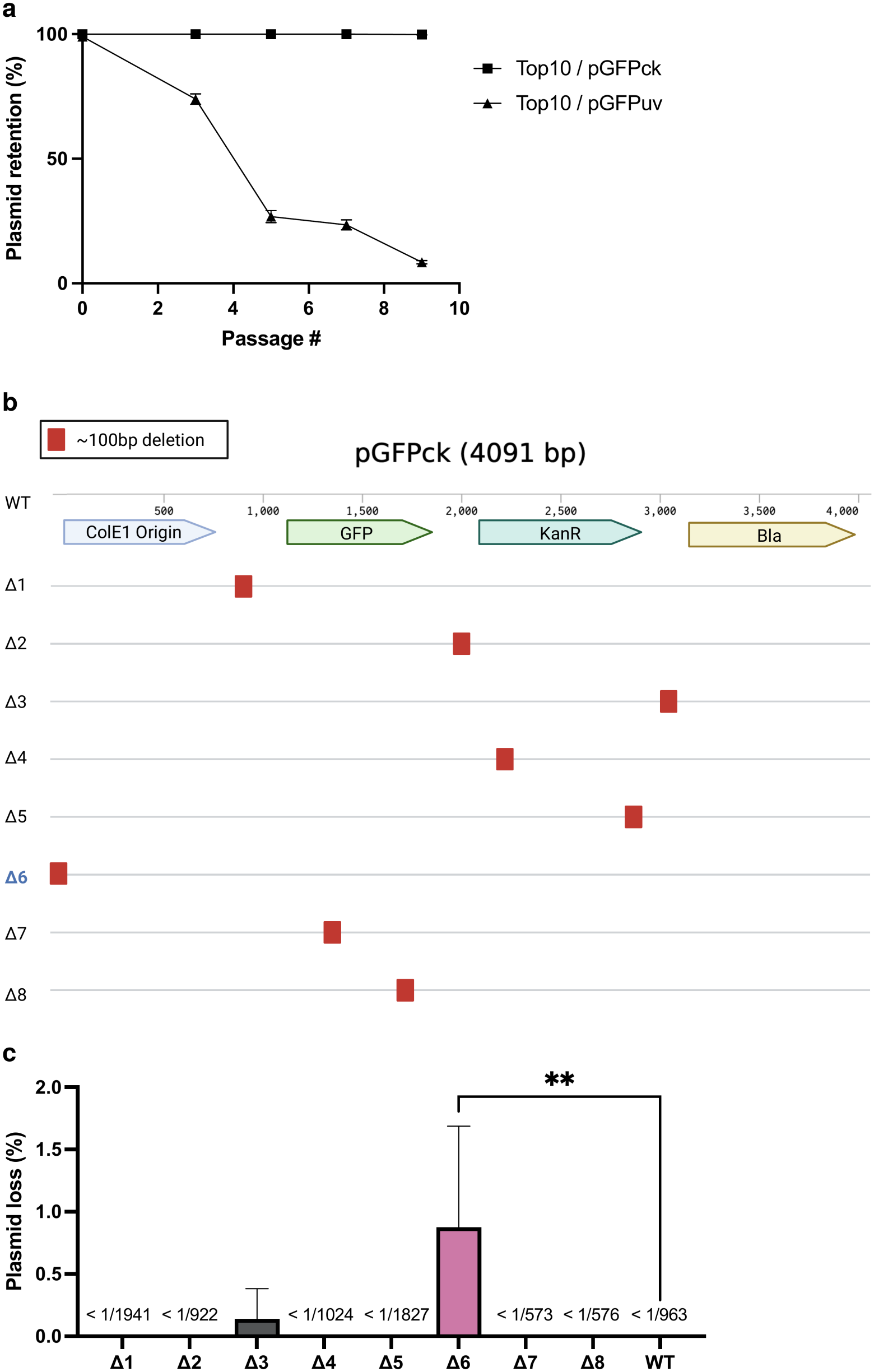
Logistics of using kanamycin as a selecting agent for early plasmid loss detection. **(a)** Plasmid stability of Top10/pGFPck (black squares) versus Top10/pGFPuv (black triangles) over several passages without selection; plasmid retention (expressed as percentage) was scored each passage as the fraction of colonies remaining fluorescent under UV. Points represent the average of a triplicate and error bars represent the standard deviation. **(b)** Map of pGFPck (4091 bp) showing the ColE1 origin, GFP, KanR, and bla (B-lac), and the location of eight ∼100 bp microdeletions (Δ1– Δ8) tiled across the sequence. **(c)** Early plasmid loss measurements (in % plasmidless colonies) for eight ∼100 bp microdeletions (Δ1–Δ8) and WT pGFPck. Loss was scored by loss of colony fluorescence for Δ1–Δ6 and WT; because the Δ7 and Δ8 deletions fall within GFP and abolish fluorescence independently of plasmid retention, those two were scored by loss of carbenicillin resistance instead. Only Δ6 (∼0.88%) differed from WT (one-way ANOVA, P = 0.0200; Dunnett’s test, adjusted P = 0.0082). Δ3 yielded a single plasmid-free colony in one of three replicates (1/1,173 colonies) and did not differ significantly from WT; all other constructs were below detection (limits shown as < 1/N).Bars represent the average of triplicate samples and error bars represent the standard deviation.

To better understand the effects of the pGFPck and the Δ6 microdeletion on plasmid stability, we measured their impacts on host fitness and on plasmid copy number (**Fig. 8 a, b**). On the fitness side, we found that the Δ6 microdeletion exhibits a higher growth rate comparable to the pGFPuv WT plasmid, reversing the reduced fitness of its parental pGFPck (**Fig. 8a**), whereas on the plasmid copy number side, we observed that the Δ6 microdeletion exhibited a substantial drop in plasmid copy number: 28.6 plasmids per chromosome, compared to 156.8, a 5.5-fold decrease.

**Figure 8.**
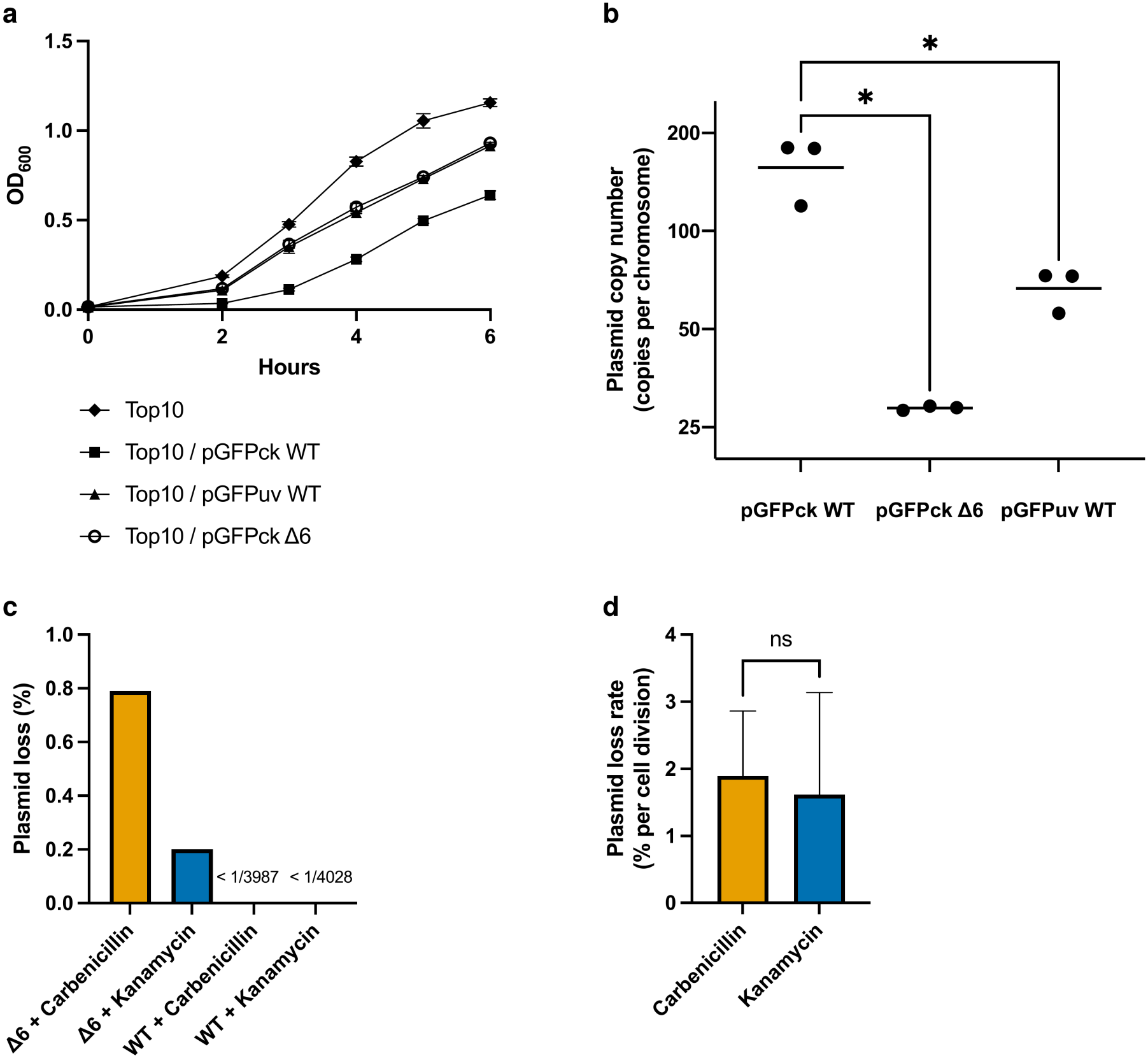
Characterization of pGFPck Δ6 and comparison of its early plasmid loss under carbenicillin and kanamycin selection. **(a)** Growth of plasmid-free Top10 (black diamond) and Top10 carrying pGFPck WT (black square), pGFPck Δ6 (open circle) or pGFPuv WT (black triangle), monitored by OD_600_ over 6 h as in **Fig. 2a**. Points are mean ± s.d., n = 3 biological replicates. **(b)** Plasmid copy number (copies per chromosome) of pGFPck WT, pGFPck Δ6 and pGFPuv WT, determined by qPCR. Points are individual biological replicates (n = 3) and horizontal lines indicate the geometric mean;*, adjusted P = 0.013 (pGFPck WT vs Δ6). pGFPck WT copy number was also higher than that of pGFPuv WT (Fig. 2b; adjusted P = 0.031). Statistical analysis as in Fig. 2b. **(c)** Early plasmid loss for pGFPck Δ6 and WT measured with each selecting agent, using loss of colony fluorescence as the readout. Values are pooled across three independent replicates; plasmid loss (%): Δ6 = 0.788% (42 dark/5329 colonies) with carbenicillin and 0.20% (9 dark/4448 colonies) with kanamycin; WT showed no detectable loss with either agent (<1/3987 carb, <1/4028 kan). Values are pooled counts rather than replicate means. **(d)** Plasmid loss rate per cell division for pGFPck Δ6, by selecting treatment. The plasmidless fractions in (c) were converted to a probability of plasmid loss per cell division using the mathematical model described in Methods, using plating efficiency data of Top10 plasmid-free cells (**Fig. 6**) and growth data corresponding to Top10/pGFPck Δ6; plasmidless generation was scored by loss of colony fluorescence as in (c). Bars are the mean of three independent biological replicates and error bars are the sample standard deviation (n = 3 biological replicates): 0.0189 ± 0.0079 for carbenicillin and 0.0162 ± 0.0124 for kanamycin, corresponding to approximately one plasmidless daughter per 53 and per 65 cell divisions respectively. Plasmid loss rates under the two drugs did not differ significantly (paired two-tailed t-test, t = 0.59, df = 2, P = 0.61).

The Δ6 microdeletion was unstable enough to allow us to carry out our comparative early plasmid loss quantification. The fraction of plasmidless cells detected in the presence of carbenicillin was 0.79%, while in the presence of kanamycin this fraction went down to 0.2%, as expected given kanamycin’s faster cytotoxicity kinetics **(Fig. 8c)**. No plasmidless colonies were detected for the more stable WT pGFPck plasmid under either selection condition, with detection limits of <1/3987 for carbenicillin and <1/4028 for kanamycin.

### Mathematical model of plasmidless cell generation

Assuming that the observed instances of plasmidless cell generation were solely due to a plasmidless daughter cell segregation and that plasmid-bearing cells do not die during replication, we built the following mathematical model to estimate the frequency of plasmidless cell generation per cell division. We used differential equations, as they allow us to consider situations where a value depends on a changing state of a given variable. In context of the method described in this paper, differential equations allow us to appropriately consider population dynamics because the total number of plasmidless cells generated at a point of time is dependent on the population state at that point of time. Our assumptions outlined in the methods section imply that change in the plasmid-bearing population with respect to time depends on a growth rate, β, multiplied by the population state, and the rate of plasmidless generation with respect to time, *L*, in proportion to population state. The change in the plasmidless population is directly the negative part of the change in the plasmid-bearing population multiplied by the probability a plasmidless cell generated at time *t* is observed upon plating at time T, *p(T-t)*, for which the viability data is used **(Fig. 6)**. This leaves us with the change in the plasmid-bearing and plasmidless populations modeled as the following differential equations,

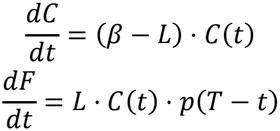

which we solve for and are implemented in the numerical solver for the loss rate as detailed in methods. Our mathematical model yielded similar plasmid loss rates per cell division of 0.0189 ± 0.0079 for carbenicillin and 0.0162 ± 0.0124 for kanamycin, (mean ± s.d., n = 3) **(Fig. 8d)**. With the sample size available, we are unable to state that the result is statistically identical, but the convergence does strongly suggest that the estimated loss rate is likely largely dependent on shared properties of the plasmid, as opposed to depending on the properties associated with the antibiotics used for the selection. Applying our mathematical model to the pGFPuv and DAS G512A early loss experiments (Fig. 2a.) produced estimates of 0.039 and <0.0011, respectively.

## Discussion

Small, multicopy plasmids are ubiquitously found in nature ^19^. Given that small, multicopy plasmids frequently lack specialized maintenance and transmission systems, their observed long-term associations with their hosts are particularly paradoxical.

Here we use a small ColE1 plasmid as a model system to estimate the frequency of spontaneous plasmidless cell generation per cell division for a ColE1-like plasmid and for a superstable mutant derivative. Our approach consists of quantifying the frequency of spontaneous, plasmidless cell generation over a short, defined time window that can be converted through a mathematical model to frequency of plasmid loss per cell division. Thus, our approach largely removes the increased fitness of plasmidless cells as a factor contributing to plasmid stability, making the study of mechanisms modulating plasmidless cell generation much more accessible.

Our method takes advantage of the transient survival of plasmid-free cells in presence of selective antibiotic. In the case of carbenicillin, the main reason for the substantial lag in killing is that the carbenicillin-induced cell wall weakening initially results in the formation of a bulge that is reversible because it can be stabilized by the outer membrane ^32,36^. It could be argued that heteroresistance ^37,38^ could underestimate the prevalence of plasmidless cells in the population, as some cells are known to stochastically enter a dormant state that protects them from the action of antibiotics, particularly those that preferentially target rapidly replicating cells such as carbenicillin ^39^. However, the strong concordance between the fluorescence and carbenicillin resistance markers rules out heteroresistance as a confounding factor, at least in our model system.

It is interesting to note that PE seems to be a more sensitive readout than vital staining, both for carbenicillin-induced and kanamycin-induced cytotoxicity. The higher sensitivity of PE relative to SYTO9/ PI staining as an indicator of cytotoxicity is a phenomenon previously reported for a variety of cytotoxic treatments in environmental bacteria ^40^ and bifidobacterium ^41^ and which suggests that cells may be committed to dying before membrane integrity is substantially altered.

Our approach for detecting the frequency of spontaneous plasmid loss in a population represents an alternative to plasmid visualization methods. These methods are challenging, as they involve introducing reporter sequence and a potentially highly labor-intensive screening. Plasmid visualization reporters typically consist of plasmid-binding proteins fused to a fluorescent protein tag, which typically requires extensive sequence engineering. Examples include the *parBC* system and fluorescence repressor-operator systems (FROS) ^42^. FROS involves an operator site and a fluorescently tagged repressor. To amplify the signal, the operator sequence is repeated in tandem several hundred times. Given that multicopy plasmids are typically small (0.8-10 kb) ^43^, and that cargo tends to destabilize them ^44^, the addition of several thousand additional basepairs of tandem operator sequence likely alters the frequency of plasmidless cell generation. Further, given that for plasmid visualization methods scoring plasmid loss relies on signal loss, plasmid loss detection depends on establishing a signal threshold with confidence. One of the strengths of our method is that it establishes plasmid loss with certainty because it produces binary outcomes.

Since no fluorescence or luminescence reporter sequence is needed, our method can be used for naturally occurring plasmids, including clinical and environmental ones. All that is required is the presence of an antibiotic resistance marker and the absence of specialized plasmid retention systems, as these mechanisms would suppress the frequency of plasmidless cell generation, confounding the interpretation of the final results. Thus, our approach opens the door for probing the range of spontaneous plasmidless cell generation frequencies found in naturally occurring multicopy plasmids.

Random distribution models predict exceedingly low frequencies of plasmid loss (< 10^-6^) for multicopy plasmids ^22^. Here we measured a copy number of ∼67 copies per chromosome for pGFPuv WT (**Fig. 3a**). If each copy segregated independently, the random distribution model would predict a frequency of plasmidless cell generation below 10^−20^, roughly eighteen orders of magnitude below the 0.034 that we measure, implying that our plasmids have a much smaller number of partitioning units or that the distribution of these plasmids is not random (**Fig. 3a**).

Further, in the work presented here, the effects of altering plasmid copy number on plasmid segregation accuracy are not consistent with a random distribution. For example, the DAS G512A and pGPFuv plasmids have comparable plasmid copy numbers, yet DAS G512A exhibits a much higher plasmid segregation accuracy (**Fig. 2b**). Also, relative to pGFPuv, the Δ6 pGFPck microdeletion exhibited a 2.3-fold drop in copy number, yet it exhibited substantially more accurate plasmid segregation (0.017 plasmid loss per cell division, compared to 0,034). These observations further reinforce the conclusions from a previous report from our group, showing that in our model system long-term plasmid stability does not correlate with plasmid copy number ^27^.

Our work implies that the mechanism underlying the observed enhancement in long-term stability associated with the DAS G512A plasmid is due to a lower frequency of plasmidless cell generation because this plasmid exhibits a comparable impact on fitness relative to its parental, unstable pGFPuv (**Fig. 2a**). To our knowledge, this represents one of the first documented examples of increased long-term plasmid stability where the main driver is not a fitness advantage of the plasmidless cells. We also report that the addition of the *nptII* gene in the pGFPck plasmid substantially increases plasmid segregation accuracy relative to the parental plasmid pGFPuv. Thus, our work establishes that plasmid segregation accuracy can be modulated by changes in sequence both in the *ori* and outside the *ori* sequence. The mechanisms involved remain unknown. In the case of pGFPck plasmid, it may have to do with increased plasmid size. It has been previously suggested that small plasmid size may be linked to instability in multicopy plasmids carrying antibiotic resistance genes ^44^, although the mechanism is not clear. Alternatively, a relative constant supercoiling *density* in vivo ^45^ (maintained at approximately σ ≈ - 0.06 by type I and II topoisomerases) means that larger plasmids have a proportionally greater number of supercoils, which in turn could affect plasmid stability ^46^. Finally, smaller plasmids may be more prone to transcriptional readthrough into the ColE1 *ori*, particularly if their cargo gene expression is driven by a strong promoter. Transcriptional readthrough into the ColE1 *ori* is known to affect replication initiation^47^ and could also affect plasmid stability. Indeed, the addition of a kanamycin resistance gene *nptII* in pGFPck substantially increases the distance between the lac promoter and the *ori*, likely decreasing transcriptional readthrough into the *ori* and possibly explaining the observed increase in plasmid copy number.

This work also explored the impact of 100 bp microdeletions on the frequency of plasmidless cell generation as an indication of how sensitive multicopy plasmid sequence is to sequence modification. Our results show remarkable robustness to microdeletions. Of the eight microdeletions that we tested, the only one with a clear destabilizing effect was a partial deletion of the 5’ end of the ColE1 *ori*. This microdeletion involves the antisense regulatory section, the same section that is duplicated in our superstable mutant (although the deletion is smaller -100 bp-relative to our superstable mutant -242 bp-), again pointing at the 5’ end of the ColE1 *ori* as an area that is important for the regulation of plasmid segregation accuracy. The Δ6 pGFPck deletion also reduced plasmid copy number ∼5.5-fold relative to pGFPck WT (**Fig. 8b**), suggesting that the removal of this portion of the ori affects replication initiation. As noted above, though, plasmid copy number does not account for plasmid segregation accuracy as the Δ6 carries fewer copies than pGFPuv WT yet exhibits more accurate plasmid segregation.

Overall, the results of pGFPck microdeletion scanning suggest that moderate sequence manipulations do not have a major effect on early plasmid loss in multicopy plasmids unless they are in the *ori* itself. This observed robustness to microdeletions is relevant for the design of plasmid tagging approaches, suggesting that small changes (<100 bp) in size do not detectably affect the frequency of plasmidless cell generation.

Our method has admittedly several potential caveats. The antibiotic itself may impact plasmidless segregation frequency by altering the distribution of plasmids within the cell or by increasing plasmid copy number. Carbenicillin, for example, is known to induce filamentation ^32^, while chloramphenicol (and potentially other inhibitors of prokaryotic translation) increase plasmid copy number ^48^. We addressed these caveats by producing consistent estimates of spontaneous plasmidless cell generation frequency using two antibiotics that have completely different mechanisms of action: carbenicillin, a beta-lactam that weakens the bacterial cell wall by inhibiting transpeptidation ^32^, and kanamycin, an aminoglycoside antibiotic that inhibits protein synthesis ^35^. Finally, our mathematical model assumes that plasmid loss is primarily the result of asymmetric partition during cell division, which is the assumption of most published mathematical models ^21,23,49^, but this is still an assumption. Plasmid loss could result from other mechanisms such as integration into the chromosome ^18,50^ or degradation ^51^.

In conclusion, we present a low-tech and efficient method to estimate the frequency of spontaneous plasmid loss during cell division assuming that inaccurate segregation is the main mechanism responsible for plasmid loss in our model system. Our method draws a clear distinction between the spontaneous frequency of plasmid loss and the increased fitness of plasmidless cells as contributors to long-term plasmid stability and is very simple, only requiring a selectable marker or a reporter, and allows the screening of large numbers of cells. Thus, our approach is ideal for the study of the stability of naturally occurring plasmids or for the identification of genetic or pharmacological modulators modulating plasmid stability. Given that small, multicopy plasmids are the platform of choice for recombinant gene expression ^5,6^, modulating plasmid stability is relevant to attempts to optimize recombinant yield ^52^.

## Methods

### Strains and plasmids

Hosts are Top10 (*F-mcrA Δ(mrr-hsdRMS-mcrBC) φ80 lacZΔM15 ΔlacX74 recA1 araD139 Δ(ara-leu)7697 galU galK rpsL (StrR) endA1 nupG*). Plasmids pGFPuv and pGFPck are described in detail in [28]. These plasmids contain pUC19 (a ColE1-like plasmid origin of replication), TEM-1 β-lactamase as a selectable marker, and cycle 3 GFP as a fluorescent reporter; pGFPck additionally carries a kanamycin resistance gene. The pGFPuv derivative DAS G512A, which contains a double mutation in the plasmid *ori, was* described in [27]. The FASTA files for all the plasmids used in this study can be found in the Supplementary Information.

### Growth conditions

Glycerol stocks of cells transformed with plasmids were prepared by mixing 600 µL of an overnight culture of the bacteria carrying the relevant plasmids to 400 µL of 50% glycerol. Glycerol stocks are stored at -80 °C. To recover the cells from the glycerol stock, frozen pellets of the glycerol stock were grown in LB media containing a final concentration of 100 µg/mL carbenicillin. Cultures were grown overnight at 37 °C with shaking at 200 rpm. For growth curves (Fig. 2a and 8a), overnight cultures were diluted to OD_600_ 0.01 in LB (Top10) or LB containing 100 µg/mL carbenicillin (plasmid-bearing Top10) and grown at 37 °C with shaking. OD_600_ was measured with a spectrophotometer at 0 h and hourly from 2 to 6 h.

### Plasmid transformation

Cells were transformed by chemical transformation. Chemically competent cells were prepared by resuspension in a 100 mM CaCl_2_ solution. Between 1 and 3 µL plasmid solution were added to the chemically competent cells, mixed and left on ice for 30 minutes. Cell-plasmid mixture was then heat-shocked at 42 °C for 1.5 minutes and transferred back on ice for five minutes. LB media was added to the cell-plasmid mixture and incubated with shaking at 37 °C for cell recovery for 30 minutes. Cells were then plated on LB agar supplemented with 100 µg/mL carbenicillin.

### Early plasmid loss assays

#### Assay based on loss of colony GFP fluorescence

Cells carrying pGFPuv (WT or DAS G512A) or pGFPck (WT or Δ1–Δ6) were grown in LB containing 100 µg/mL carbenicillin, or 50 µg/mL kanamycin for the comparison in Fig. 8c, to OD_600_ 0.4, diluted ∼5 × 10^−6^ and plated on LB agar without antibiotic. Colonies that grew after an overnight incubation at 37 °C were observed under UV lighting. Individual GFP^lo^ colonies on the LB agar plate were indicative of individual plasmid loss events. The results were expressed as the fraction of GFP^lo^ colonies relative to the total colony number, in percentage. This method is illustrated in **Fig. 1**.

#### Assay based on the loss of carbenicillin resistance in liquid

Cells transformed with a given plasmid (pGFPuv WT or DAS G512A; pGFPck Δ7 and Δ8, Fig. 7c) were grown in liquid culture in the presence of carbenicillin to OD_600_ 0.4 and diluted for plating on LB agar without carbenicillin. After growth on the LB agar plate, a random colony was picked and suspended in 50 µL PBS. 5 µL of this culture suspension were added to one well of a 96-well plate containing 200 µL LB. Another 5 µL of the same suspension was also added to the corresponding well of another 96-well plate containing 200 µL LB with carbenicillin. This process was repeated for all the remaining colonies on the LB agar plate. Carbenicillin-free and carbenicillin-containing 96-well plates were incubated at 37 °C overnight. Following the overnight incubation, the OD_600_ was measured using a plate reader to determine viability in the presence and absence of carbenicillin. Colonies that grew in both the presence and the absence of carbenicillin were scored as plasmid-positive. Colonies that only grew in the absence of carbenicillin were scored as plasmid-negative. This method is illustrated in **Fig. 1**.

#### Long-term plasmid retention

Top10 cells carrying pGFPuv or pGFPck were grown overnight in LB containing 100 µg/mL carbenicillin (passage 0) and then passaged without selection by diluting 1:1,000 into LB and growing for 22 h at 37 °C with shaking. At the indicated passages, cultures were serially diluted and plated on LB agar without antibiotic, and retention was scored as the percentage of colonies fluorescent under UV.

#### Plasmid copy number quantification by qPCR

Plasmid copy number was quantified as the ratio of a plasmid-borne to a single-copy chromosomal amplicon in the same total-DNA sample. The plasmid target was a 105 bp amplicon in *bla* (5′-AAATCTGGAGCCGGTGAGCGTG-3′ / 5′-TGCCTGACTCCCCGTCGTGTAG-3′), sequence-identical in all constructs. The chromosomal reference was a 105 bp terminus-proximal amplicon (“*ter*”; 5′-TACTCCACTGCGCCAAATCGCC-3′ / 5′-GTCATGTCGGAGGGCACGAACG-3′) at positions 1,681,047– 1,681,151 of the sequenced *E. coli* Top10 genome, chosen so that reference dosage is insensitive to growth rate.

Three independent colonies per construct were grown overnight at 37 °C in LB containing 100 µg/mL carbenicillin, subcultured 1:100 into the same medium, and harvested in exponential phase at OD_600_ 0.4. Total DNA was extracted with the DNeasy Blood and Tissue kit (QIAGEN, Gram-negative pretreatment) and quantified by Qubit dsDNA assay. Equal masses (750 ng) were digested with HindIII, which has no site within either amplicon. Digestion was confirmed by agarose gel.

Reactions (20 µL) contained 1× SYBR Green master mix, 400 nM each primer and 2 ng digested DNA, with the two primer pairs in separate wells. Cycling on a Bio-Rad CFX96 was 95 °C for 3 min, then 40 cycles of 95 °C for 10 s and 60 °C for 30 s, followed by melt-curve analysis; both amplicons gave a single melt peak. Amplification efficiency determined from a five-point ten-fold dilution series in technical triplicate was 93.3% for *bla* (R^2^ 0.997) and 96.4% for *ter* (R^2^ 0.995), and copy number was calculated as 2^(Cq_*ter* − Cq_*bla*). Samples on separate plates were bridged using two samples common to both. No-template controls gave no amplification, and total DNA from untransformed Top10 gave normal *ter* amplification with no *bla* amplification.

Samples were run in technical duplicate per amplicon, averaged to one Cq per sample per amplicon, and rejected if replicates spanned more than 0.6 Cq. Copy numbers are reported as plasmid copies per chromosome from n = 3 biological replicates, as geometric means with individual replicates shown. Groups were compared by Welch’s one-way ANOVA assuming a lognormal distribution, followed by Dunnett’s T3 multiple comparisons test (pGFPuv WT vs DAS G512A, pGFPck WT vs Δ6, and pGFPuv WT vs pGFPck WT).

#### Kinetics of cell death in the presence of antibiotics as determined by plating efficiency

Top10 cells, plasmid-free or carrying pGFPck as a resistant control (Fig. 4b), were grown in 25 mL LB without antibiotics and incubated at 37 °C shaking at 200 rpm. At OD_600_ of 0.4, antibiotics were added to the culture (none for the untreated control), and incubation was resumed at 37 °C, still shaking at 200 rpm. The final concentrations were 100 µg/mL (carbenicillin) and 50 µg/mL (kanamycin), which correspond to standard selective concentrations on LB agar plates. Before exposure of carbenicillin or kanamycin (referred to as time 0), and at various time points during exposure, OD_600_ was measured (Fig. 4) and serial dilutions were performed for plating on LB agar without antibiotics to detect colony forming units. LB agar plates were incubated at 37 °C overnight. Plates were imaged for colony count and the result was expressed as a fraction of drug-exposed cells able to form colonies relative to untreated control.

#### Kinetics of cell death in the presence of antibiotics using vital staining

Plasmid-free Top10 cells were grown in 25 mL LB without antibiotics and incubated at 37 °C with shaking. When the culture reached an OD600 of 0.4, antibiotics were added (none for the untreated control), and incubation was resumed at 37 °C with shaking. The antibiotic concentrations that we tested corresponded to the concentrations standard for selective plates: 100 µg/mL or 50 µg/mL (carbenicillin and kanamycin, respectively). Before exposure of carbenicillin or kanamycin (time 0), and at various time points during exposure, samples were prepared for staining using LIVE/DEAD− BacLight− Bacterial Viability Kit. For staining, 3 mL of the culture was centrifuged at 11,000 rpm for four minutes. The pellet was washed twice with 1.5 mL 0.85% NaCl using centrifugation at 3,000 rpm for five minutes during each wash. The pellet was resuspended in 900 µL 0.85% NaCl. Working solutions of propidium iodide (PI) and SYTO 9 were prepared to final concentrations 400 µM PI and 33.4 µM SYTO 9. 50 µL of each working solution were added to the 900 µL NaCl-resuspension. The mixture was incubated in the dark for 15 minutes. Stained samples were imaged on a Solamere spinning-disk confocal microscope with a 63× oil-immersion objective, using the GFP channel for SYTO 9 and the RFP channel for PI, with the same acquisition settings for all samples. Brightness and contrast were adjusted linearly in Fiji and applied identically to all images within each channel. Cells were scored manually as live (SYTO 9 only) or dead (PI-positive) in one field per biological replicate.

#### Generation of microdeletion constructs

Eight ∼100 bp microdeletions spanning the pGFPck plasmid were generated using the Q5^®^ Site-Directed Mutagenesis Kit. Deletion primers were designed using the NEBaseChanger tool (deletion positions shown in **Fig. 7b**; primers and coordinates in **Table S1**). Mutagenesis was performed according to the manufacturer’s instructions and transformed into chemically competent Top10 cells. Transformants were selected on LB agar carbenicillin (100 µg/mL), and plasmids from individual colonies were verified by Sanger sequencing.

#### Model of plasmidless generation by asymmetrical cell division

The plating assays return the fraction of colonies that are plasmidless after growth under antibiotic selection. We model the behavior of plasmid containing bacteria and plasmidless bacteria over the course of exposure to a selected antibiotic using a system of ordinary differential equations that confers a single unknown variable, the loss rate occurring by asymmetrical cell division, in units of probability per division. From the system, the plasmid loss rate corresponding to a set of data is solved for numerically.

There are three inputs into this model:

1. The growth curve of the strain of bacteria with plasmid, from which a doubling time is directly inferred from the OD600 data (*Growth conditions*).
2. The survival rates with respect to time of exposure of bacteria normalized as a proportion (*Kinetics of cell death in the presence of antibiotics as determined by plating efficiency*).
3. The final observed counts of colonies that express the plasmid gene and colonies which have lost the plasmid (*Assay based on loss of colony GFP fluorescence*).

For each biological replicate, colony counts from technical replicate plates were averaged before use in computations.

For the description of the model we define the following symbols:

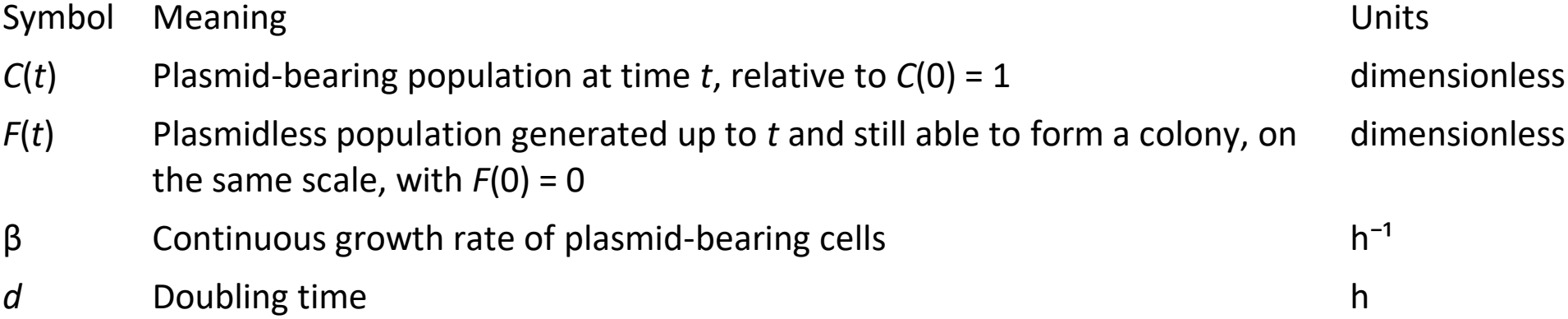
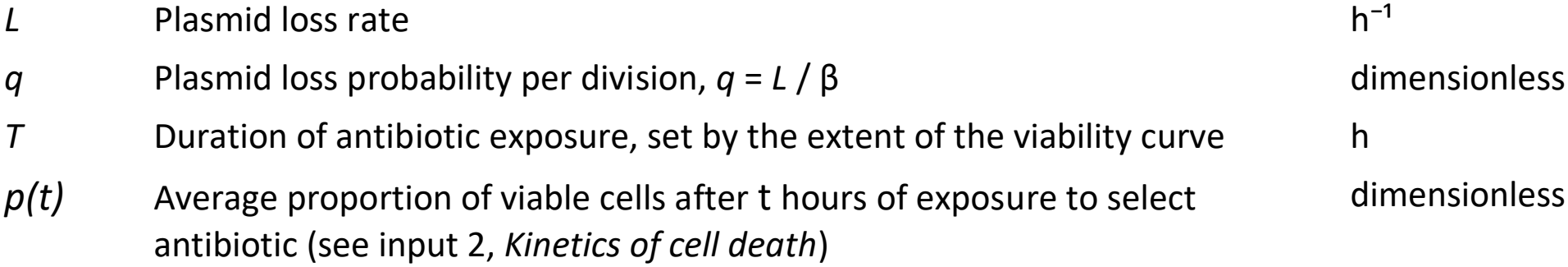

| Symbol | Meaning | Units |
|---|---|---|
| *C*(*t*) | Plasmid-bearing population at time *t*, relative to *C*(0) = 1 | dimensionless |
| *F*(*t*) | Plasmidless population generated up to *t* and still able to form a colony, on the same scale, with *F*(0) = 0 | dimensionless |
| β | Continuous growth rate of plasmid-bearing cells | h<sup>−1</sup> |
| *d* | Doubling time | h |
| *L* | Plasmid loss rate | h<sup>−1</sup> |
| *q* | Plasmid loss probability per division, *q* = *L* / β | dimensionless |
| *T* | Duration of antibiotic exposure, set by the extent of the viability curve | h |
| *p(t)* | Average proportion of viable cells after t hours of exposure to select antibiotic (see input 2, *Kinetics of cell death*) | dimensionless |

For the model we need to make a series of assumptions to arrive at the initial set of differential equations, we list ones made as follows:

1. Plasmid loss occurs at division, when a daughter cell inherits no plasmid.
2. The loss probability is constant over the course of antibiotic exposure, and independent of culture density and time of exposure to antibiotic.
3. Plasmid bearing cells do not die over the course of antibiotic exposure.
4. Plasmidless cells do not replicate after generation. This is an unnecessary assumption for Kanamycin exposure, since T_kanamycin_ < *d*, but for carbenicillin this may influence computations.
5. Loss events are rare relative to total divisions, which allows us to use the growth curve data we generated for estimation of *d* as is.
6. We assume that a linear interpolation of the survival curve data points is satisfactory. In reality, a linear interpolation is not a true representation of the data, with error increasing with increasing time interval lengths.
7. Plasmidless cells generated by segregation have the same survival function as the plasmidless cells used to measure it. In reality, a plasmidless daughter cell may inherit some pool of the protein encoded for by the antibiotic resistance cassette which could increase its rate of survival.
8. OD_600_ is proportional to cell count and we can use OD_600_ measurements to directly infer a doubling time.
9. We only consider the interval [0, *T*] for *t* and a time value where < 0.1% cells are viable on average is sufficient to consider for the value of *T*.

### Computation of doubling time

We compute doubling time in this experiment by computing doubling time for every sequential interval from the growth curve data and select *d* such that it is computed from the interval which minimizes *d*. This is intended to represent doubling time corresponding to an exponential growth phase, which matches conditions used in observing generation of plasmidless cells.

#### Derivation of key formulas

From the assumptions, we arrive at the following system of differential equations:

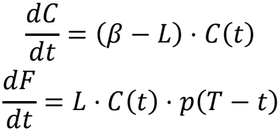

Which when solved individually for *C(t)* and *F(t)*, give the following equations:

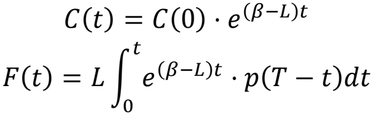

In context of our experiment, we can set *C*(0) = 1, since it represents 100% of the population at t = 0. Note that we cannot find the antiderivative of the expression for F(t) analytically due to the nature of the viability data. However, it should be noted that the initial condition is additive and is omitted as *F*(0) represents none of the population at t = 0. At time T, *F*(*T*) represents the proportion of plasmidless colonies relative to C(0), and C(T) represents the proportion of plasmid bearing colonies relative to *C*(0). Therefore, the number of plasmidless colonies divided by the number of plasmid-bearing colonies observed at plating (see input 3.) gives us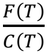 the following equation with only one unknown, L., which gives us

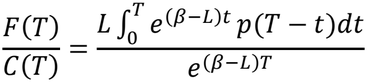

The function, *p*(*T* − *t*), is linearly interpolated with the numPy function, interp, and integrated with the integrate.quad function from the sciPy library. We then arrange the equation so that it equals zero and numerically find the value of L that satisfies the equation using Brent’s method (brentq) as a root finder.

#### Solving for β

β is the continuous growth rate with respect to time. We can use the continuous growth equation, 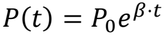, to solve for β, where P is the population size relative to *P*_0_ at time t and *P*_0_ is the population size at *t* = 0. Specializing the equation with the doubling time, *d, P*(0) = 1, and *P*(*d*) = 2 gives us 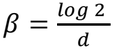, which we subsequently use for computations in our model’s implementation.

#### Conversion to loss probability per division

The final output of the model, *q*, is in units of loss probability per division, which is given by dividing the total number of plasmids lost, 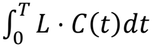, by the total number of divisions, 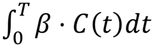. The integrals cancel after factoring out constants *L* and *β*, leaving us with 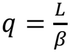.

## Supporting information

Plasmid Sequences

## Data availability

The code, computational environment and data used for the mathematical model are available at https://github.com/vj932/script_data_pl. FASTA files for all plasmids used in this study are provided in the Supplementary Information.

## Acknowledgements

CE was supported by a Graduate Fellowship for Innovators from UCSC’s California Institute for Quantitative Biosciences (QB3). The authors would also like to thank Ryan Alex for help building the mathematical model in the early stages of the manuscript .

## Author contributions

CE conducted all the experimental work, contributed to the design of the project, to the writing of the manuscript, and formatted the manuscript for publication. V.A. built the mathematical model and wrote the corresponding methods, results and discussion sections of the article. MC contributed to the overall design of the project, to the writing of the manuscript, and conducted the final edits.

## Additional information

### Supplementary Information

accompanies this paper (Supplementary Table S1; plasmid FASTA files).

## References

1 Smalla, K., Jechalke, S. & Top, E. M. Plasmid Detection, Characterization, and Ecology. Microbiol Spectr 3, PLAS–0038–2014 (2015). 10.1128/microbiolspec.PLAS-0038-2014

2 Werren, J. H. Selfish genetic elements, genetic conflict, and evolutionary innovation. Proc Natl Acad Sci U S A 108 Suppl 2, 10863–10870 (2011). 10.1073/pnas.1102343108

3 Anda, M. et al. Bacterial clade with the ribosomal RNA operon on a small plasmid rather than the chromosome. Proc Natl Acad Sci U S A 112, 14343–14347 (2015). 10.1073/pnas.1514326112

4 Castaneda-Barba, S., Top, E. M. & Stalder, T. Plasmids, a molecular cornerstone of antimicrobial resistance in the One Health era. Nat Rev Microbiol 22, 18–32 (2024). 10.1038/s41579-023-00926-x

5 Million-Weaver, S., Alexander, D. L., Allen, J. M. & Camps, M. Quantifying plasmid copy number to investigate plasmid dosage effects associated with directed protein evolution. Methods Mol Biol 834, 33–48 (2012). 10.1007/978-1-61779-483-4_3

6 Camps, M. Modulation of ColE1-like plasmid replication for recombinant gene expression. Recent Pat DNA Gene Seq 4, 58–73 (2010). 10.2174/187221510790410822

7 Baltrus, D. A. Exploring the costs of horizontal gene transfer. Trends Ecol Evol 28, 489–495 (2013). 10.1016/j.tree.2013.04.002

8 Porse, A., Schonning, K., Munck, C. & Sommer, M. O. Survival and Evolution of a Large Multidrug Resistance Plasmid in New Clinical Bacterial Hosts. Mol Biol Evol 33, 2860–2873 (2016). 10.1093/molbev/msw163

9 San Millan, A. & MacLean, R. C. Fitness Costs of Plasmids: a Limit to Plasmid Transmission. Microbiol Spectr 5 (2017). 10.1128/microbiolspec.MTBP-0016-2017

10 San Millan, A. et al. Integrative analysis of fitness and metabolic effects of plasmids in Pseudomonas aeruginosa PAO1. ISME J 12, 3014–3024 (2018). 10.1038/s41396-018-0224-8

11 Harrison, E. & Brockhurst, M. A. Plasmid-mediated horizontal gene transfer is a coevolutionary process. Trends Microbiol 20, 262–267 (2012). 10.1016/j.tim.2012.04.003

12 Rankin, D. J., Rocha, E. P. & Brown, S. P. What traits are carried on mobile genetic elements, and why? Heredity (Edinb) 106, 1–10 (2011). 10.1038/hdy.2010.24

13 Hall, J. P. J. et al. Plasmid fitness costs are caused by specific genetic conflicts enabling resolution by compensatory mutation. PLoS Biol 19, e3001225 (2021). 10.1371/journal.pbio.3001225

14 Harms, A., Brodersen, D. E., Mitarai, N. & Gerdes, K. Toxins, Targets, and Triggers: An Overview of Toxin-Antitoxin Biology. Mol Cell 70, 768–784 (2018). 10.1016/j.molcel.2018.01.003

15 Gerdes, K., Howard, M. & Szardenings, F. Pushing and pulling in prokaryotic DNA segregation. Cell 141, 927–942 (2010). 10.1016/j.cell.2010.05.033

16 Bouet, J. Y. & Funnell, B. E. Plasmid Localization and Partition in Enterobacteriaceae. EcoSal Plus 8 (2019). 10.1128/ecosalplus.ESP-0003-2019

17 Brockhurst, M. A. & Harrison, E. Ecological and evolutionary solutions to the plasmid paradox. Trends Microbiol 30, 534–543 (2022). 10.1016/j.tim.2021.11.001

18 Carroll, A. C. & Wong, A. Plasmid persistence: costs, benefits, and the plasmid paradox. Can J Microbiol 64, 293–304 (2018). 10.1139/cjm-2017-0609

19 Finks, S. S. & Martiny, J. B. H. Plasmid-Encoded Traits Vary across Environments. mBio 14, e0319122 (2023). 10.1128/mbio.03191-22

20 Stoesser, N. et al. Evolutionary History of the Global Emergence of the Escherichia coli Epidemic Clone ST131. mBio 7, e02162 (2016). 10.1128/mBio.02162-15

21 Cooper, N. S., Brown, M. E. & Caulcott, C. A. A mathematical method for analysing plasmid stability in micro-organisms. J Gen Microbiol 133, 1871–1880 (1987). 10.1099/00221287-133-7-1871

22 De Gelder, L. et al. Combining mathematical models and statistical methods to understand and predict the dynamics of antibiotic-sensitive mutants in a population of resistant bacteria during experimental evolution. Genetics 168, 1131–1144 (2004). 10.1534/genetics.104.033431

23 Lau, B. T., Malkus, P. & Paulsson, J. New quantitative methods for measuring plasmid loss rates reveal unexpected stability. Plasmid 70, 353–361 (2013). 10.1016/j.plasmid.2013.07.007

24 Summers, D. K. & Sherratt, D. J. Multimerization of high copy number plasmids causes instability: CoIE1 encodes a determinant essential for plasmid monomerization and stability. Cell 36, 1097–1103 (1984). 10.1016/0092-8674(84)90060-6

25 Pogliano, J., Ho, T. Q., Zhong, Z. & Helinski, D. R. Multicopy plasmids are clustered and localized in Escherichia coli. Proc Natl Acad Sci U S A 98, 4486–4491 (2001). 10.1073/pnas.081075798

26 Reyes-Lamothe, R. et al. High-copy bacterial plasmids diffuse in the nucleoid-free space, replicate stochastically and are randomly partitioned at cell division. Nucleic Acids Res 42, 1042–1051 (2014). 10.1093/nar/gkt918

27 Standley, M. S., Million-Weaver, S., Alexander, D. L., Hu, S. & Camps, M. Genetic control of ColE1 plasmid stability that is independent of plasmid copy number regulation. Curr Genet 65, 179–192 (2019). 10.1007/s00294-018-0858-0

28 Standley, M., Allen, J., Cervantes, L., Lilly, J. & Camps, M. Fluorescence-Based Reporters for Detection of Mutagenesis in E. coli. Methods Enzymol 591, 159–186 (2017). 10.1016/bs.mie.2017.03.013

29 Renzette, N. et al. Localization of RecA in Escherichia coli K-12 using RecA-GFP. Mol Microbiol 57, 1074–1085 (2005). 10.1111/j.1365-2958.2005.04755.x

30 McGrew, D. A. & Knight, K. L. Molecular design and functional organization of the RecA protein. Crit Rev Biochem Mol Biol 38, 385–432 (2003). 10.1080/10409230390242489

31 Field, C. M. & Summers, D. K. Multicopy plasmid stability: revisiting the dimer catastrophe. J Theor Biol 291, 119–127 (2011). 10.1016/j.jtbi.2011.09.006

32 Yao, Z., Kahne, D. & Kishony, R. Distinct single-cell morphological dynamics under beta-lactam antibiotics. Mol Cell 48, 705–712 (2012). 10.1016/j.molcel.2012.09.016

33 Rolinson, G. N. Effect of beta-lactam antibiotics on bacterial cell growth rate. J Gen Microbiol 120, 317–323 (1980). 10.1099/00221287-120-2-317

34 Brook, I. The role of beta-lactamase-producing-bacteria in mixed infections. BMC Infect Dis 9, 202 (2009). 10.1186/1471-2334-9-202

35 Garneau-Tsodikova, S. & Labby, K. J. Mechanisms of Resistance to Aminoglycoside Antibiotics: Overview and Perspectives. Medchemcomm 7, 11–27 (2016). 10.1039/C5MD00344J

36 Daly, K. E., Huang, K. C., Wingreen, N. S. & Mukhopadhyay, R. Mechanics of membrane bulging during cell-wall disruption in gram-negative bacteria. Phys Rev E Stat Nonlin Soft Matter Phys 83, 041922 (2011). 10.1103/PhysRevE.83.041922

37 Andersson, D. I., Nicoloff, H. & Hjort, K. Mechanisms and clinical relevance of bacterial heteroresistance. Nat Rev Microbiol 17, 479–496 (2019). 10.1038/s41579-019-0218-1

38 Medaney, F., Dimitriu, T., Ellis, R. J. & Raymond, B. Live to cheat another day: bacterial dormancy facilitates the social exploitation of beta-lactamases. ISME J 10, 778–787 (2016). 10.1038/ismej.2015.154

39 Lee, A. J. et al. Robust, linear correlations between growth rates and beta-lactam-mediated lysis rates. Proc Natl Acad Sci U S A 115, 4069–4074 (2018). 10.1073/pnas.1719504115

40 Shi, L. et al. Limits of propidium iodide as a cell viability indicator for environmental bacteria. Cytometry A 71, 592–598 (2007). 10.1002/cyto.a.20402

41 Amor, K. B. et al. Multiparametric flow cytometry and cell sorting for the assessment of viable, injured, and dead bifidobacterium cells during bile salt stress. Appl Environ Microbiol 68, 5209–5216 (2002). 10.1128/AEM.68.11.5209-5216.2002

42 Le, T. B. & Laub, M. T. New approaches to understanding the spatial organization of bacterial genomes. Curr Opin Microbiol 22, 15–21 (2014). 10.1016/j.mib.2014.09.014

43 Rodriguez-Beltran, J., DelaFuente, J., Leon-Sampedro, R., MacLean, R. C. & San Millan, A. Beyond horizontal gene transfer: the role of plasmids in bacterial evolution. Nat Rev Microbiol 19, 347–359 (2021). 10.1038/s41579-020-00497-1

44 Wein, T., Wang, Y., Hulter, N. F., Hammerschmidt, K. & Dagan, T. Antibiotics Interfere with the Evolution of Plasmid Stability. Curr Biol 30, 3841–3847 e3844 (2020). 10.1016/j.cub.2020.07.019

45 Kouzine, F., Sanford, S., Elisha-Feil, Z. & Levens, D. The functional response of upstream DNA to dynamic supercoiling in vivo. Nat Struct Mol Biol 15, 146–154 (2008). 10.1038/nsmb.1372

46 Singh, G., Hong, Y., Inman, J. T., Sethna, J. P. & Wang, M. D. Torsional Mechanics of Circular DNA. bioRxiv (2024).

47 Stueber, D. & Bujard, H. Transcription from efficient promoters can interfere with plasmid replication and diminish expression of plasmid specified genes. EMBO J 1, 1399–1404 (1982). 10.1002/j.1460-2075.1982.tb01329.x

48 Frenkel, L. & Bremer, H. Increased amplification of plasmids pBR322 and pBR327 by low concentrations of chloramphenicol. DNA 5, 539–544 (1986). 10.1089/dna.1.1986.5.539

49 Werbowy, O., Werbowy, S. & Kaczorowski, T. Plasmid stability analysis based on a new theoretical model employing stochastic simulations. PLoS One 12, e0183512 (2017). 10.1371/journal.pone.0183512

50 Gomberg, A. F. & Grossman, A. D. It’s complicated: relationships between integrative and conjugative elements and their bacterial hosts. Curr Opin Microbiol 82, 102556 (2024). 10.1016/j.mib.2024.102556

51 Spengler, G. et al. The mechanism of plasmid curing in bacteria. Curr Drug Targets 7, 823–841 (2006). 10.2174/138945006777709601

52 Pouresmaeil, M. & Azizi-Dargahlou, S. Factors involved in heterologous expression of proteins in E. coli host. Arch Microbiol 205, 212 (2023). 10.1007/s00203-023-03541-9

