## Supplementary Table 1 for "Efficient Method for Estimating the Frequency of Plasmidless Cell Generation per Cell Division"

**Table S1: Microdeletion primers and deletion coordinates**

| Mutant | Deleted Region (bp) | Forward Primer | Reverse Primer |
|---|---|---|---|
| Δ1 | 659:758 | GAGGAAGCGGAAGAG | CAGGAAAGAACATGTGAG |
| Δ2 | 1884:1983 | TAATACAAGGGGTGTTATG | GATCCGATATCATTTAAATCTAGG |
| Δ3 | 2830:2929 | TCAGGTGGCACTTTTC | AATTAACCAATTCTGATTAGAAAAAC |
| Δ4 | 2129:2228 | AATTTATGCCTCTTCCG | TACAATCGATAGATTGTCG |
| Δ5 | 2715:2814 | ATCAGAATTGGTTAATTGGTTG | AAACTCACCGAGGCAG |
| Δ6 | 3923:4014 | TCTCATGACCAAAATCC | AGTTACCAATGCTTAATCAG |
| Δ7 | 1062:1161 | GGTGATGCAACATACG | TCATTTTTTCTACCGGTAC |
| Δ8 | 1512:1611 | AATACTCCAATTGGCG | TGTATACATTGTGTGAGTTATAG |
